# Bifurcated Hydrogen Bonds and the Fold Switching of Lymphotactin

**DOI:** 10.1101/2020.05.11.089227

**Authors:** Prabir Khatua, Alan J Ray, Ulrich H. E. Hansmann

## Abstract

Lymphotactin (Ltn) exists under physiological conditions in an equilibrium between two interconverting structures with distinct biological functions. Using Replica-Exchange-with-Tunneling we study the conversion between the two folds. Unlike previously proposed, we find that the fold switching does not require unfolding of Lymphotactin, but proceeds through a series of intermediates that remain partially structured. This process relies on two bifurcated hydrogen bonds that connect the *β*_2_ and *β*_3_ strands and eases the transition between the hydrogen bond pattern by which the central three-stranded *β*-sheet in the two forms differ.

## 1 INTRODUCTION

Exhibiting a diverse spectrum of functions, reaching from transport of molecules to catalysis of biochemical reactions, proteins play a crucial role in the molecular machinery of cells. Protein function is determined by the three-dimensional structure. In the now classical model of protein folding the sequence of amino acids encodes an energy landscape that funnels folding pathways into a unique native state. ^1,2^ While this mechanism describes folding of many proteins, it needs to be modified for intrinsically disordered^3,4^ or metamorphic proteins^5,6^ where the sequence encodes not only a single native fold but an ensemble of structures, allowing a single protein to take multiple functions. Hence, for understanding the role of intrinsically disordered and metamorphic proteins in cells, and the regulation of their function, it is necessary to establish the underlying multi-funnel energy landscape that leads to this ensemble of diverse structures, and to comprehend the mechanism by which these structures convert into each other.

This task can be most easily tackled for metamorphic proteins such as the 93-residue protein Lymphotactin (Ltn) which are observed in two well-defined structures. As a chemokine Ltn belongs to a family of signaling proteins whose primary function is to direct immune response leukocytes toward areas of inflammation.^7,8^ Ltn has only one of the two disulfide bonds otherwise found in chemokines, allowing it to adopt and switch between two well-defined native folds with distinct functions. The first one (Ltn10) is a typical chemokine-like fold with a three-stranded *β*-sheet attached to a C-terminal *α*-helix (PDB ID: 2HDM;^9^ see Figure 1(a)). When in its second form, Ltn40, the protein forms dimers and has a four-stranded *β*-sheet in a dimeric *β*-sandwich (PDB ID: 2JP1;^10^ see Figure 1(b)). The two forms have distinct and complementary functions: Ltn10 activates the XCR1 receptor on the cell surface, while Ltn40 binds to heparin, a polysaccharide component of the extracellular matrix. ^5,11^ By being able to assume both motifs and perform disparate and complementing functions, Lymphotactin bridges the two main functional aspects of chemokine physiology: activation of specific GPCRs leading to chemotaxis,^12^ and establishing a signaling gradient toward the target location by binding with extracellular matrix GAGs. Under physiological conditions, both forms rapidly interconvert and are equally populated. However, the presence of several basic amino acids (nine Arg and six Lys residues) makes Lymphotactin sensitive to solution conditions and temperature. For example, Ltn10 is the dominant conformation at lower temperature (10°C) and high salt concentration (200 mM NaCl), while the alternative fold Ltn40 is dominant at 40°C and no salt. This shift of the equilibrium with temperature and ionic strength was investigated in CHARMM simulations in ^13,14^ which showed an accumulation of sodium ions around the charged residues of the helical region that increased with lowering temperature. These computational results are in agreement with experimental observations that high salt concentration and low temperature stabilize the chemokine fold. ^15^

**Figure 1:**
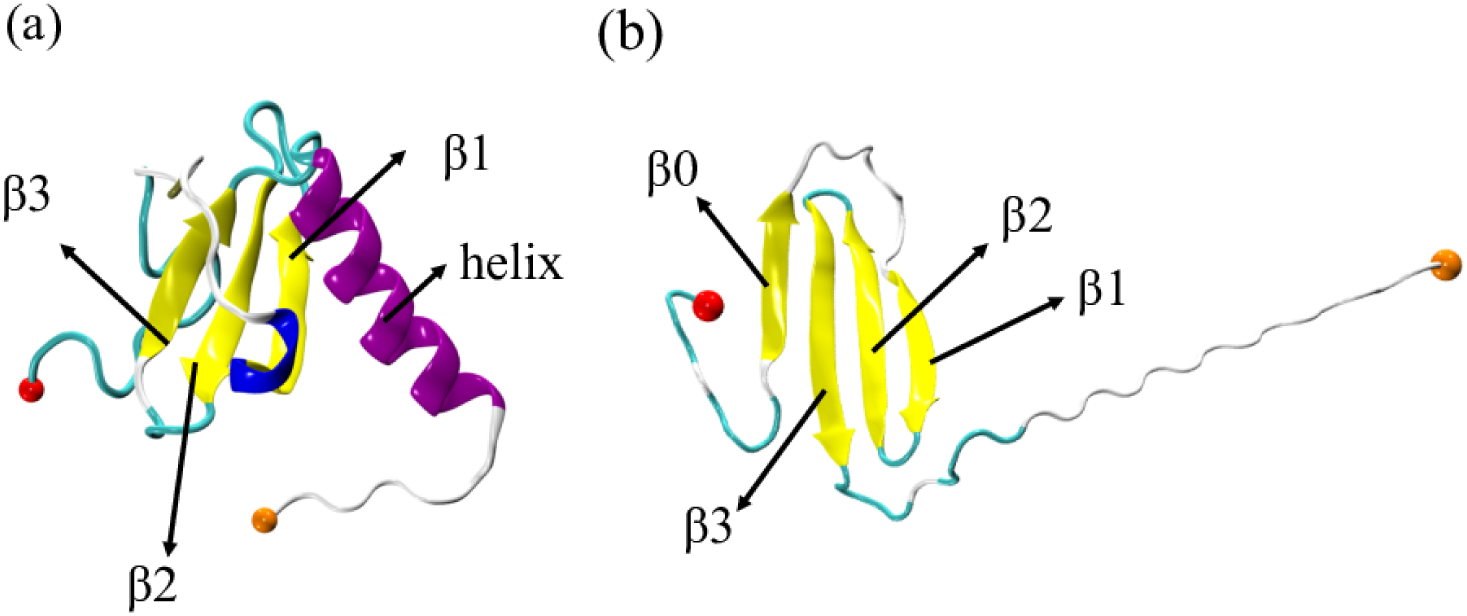
Lymphotactin chains can take two distinct structures: as deposited in the Protein Data Bank: (a) Ltn10 (PDB-ID: 2HDM) and (b) Ltn40 (PDB-ID: 2JP1). Labels identify the secondary structural elements, and the N-terminal and C-terminal C_*α*_ atoms are shown as spheres in red and orange colors, respectively.

While the structure and function of the two Lymphotactin motifs have been studied in detail,^9–11,15,16^ the mechanism of interconversion is still unclear. Unlike other metamorphic proteins, the fold switch requires a complete reorganization of core residues^5^ making the conversion especially difficult to study in experiments. Volkman’s group ^17^ examined the kinetic rates of the process by stopped-flow fluorescence. Their results suggest that the conversion process involves large-scale unfolding with a disruption of all stabilizing hydrogen bonds. However, such a mechanism is difficult to reconcile with the surprisingly low barrier separating Ltn10 and Ltn40 that rather suggests a conversion pathway going through intermediates with conserved local contacts (such as in three *β*-strands *β*_1_, *β*_2_ and *β*_3_), found in both folds and only encountering minimal disruptions of bonds. Unfortunately, such crucial but transient intermediates are hard to resolve on the short-time scales by which Ltn10 and Ltn40 convert into each other, and may have been below the temporal resolution of the Volkman’s experiments.^17^ On the other hand, the interconversion time scales are still too long to be studied with sufficient statistics in all-atom molecular dynamics simulations. While intermediates were observed in a previous computational study by Camilloni and Ludovico, ^18^ that investigation relied on a two-funneled Go-model, and it is not clear how far the presence of the intermediates and the heights of the barrier reflect the details of the construction of this model rather than the physics of the system.

In this study, we aim to resolve the experimental discrepancy by studying the Lymphotactin conversion process in all-atom simulations relying on a physical force field. As it is difficult to obtain from regular molecular dynamics sufficient statistics, we utilize an enhanced sampling technique developed in our lab, that was designed specifically for the investigation of such switching processes. Our technique, Replica-Exchange-with-Tunneling (RET),^19–22^ allows us to observe the interconversion process with sufficient detail to characterize important intermediates. This in turn enables us to propose a conversion mechanism that is consistent with the experimentally observed low barrier separating Ltn10 and Ltn40. While the experimentally observed equilibrium is between Ltn10 monomers and Ltn40 homodimers, dimerization and conversion are separate processes, with the transition between Ltn10 monomers and Ltn40 monomers being the rate-limiting process.^17^ For this reason, we consider only the conversion of monomers, but one should keep in mind that the Le Chatelier’s principle would imply a shift of the equilibrium between Ltn10 and Ltn40 monomers toward the Ltn40 form if also the subsequent dimerization was considered. Our analysis indicates that the fold switch in Lymphotactin monomers happens along a series of only partially unfolded intermediates, with the breaking and reformation of secondary structure relying on the presence of two bifurcated backbone hydrogen bonds that connect the *β*_2_ and *β*_3_ strands found in both motifs. We conjecture that these bifurcated hydrogen bonds are essential for fold switching, as they allow a repositioning of the *β*-strand forming residues without the need to cross high energy barriers.

## 2 MATERIALS AND METHODS

### 2.1 Replica-Exchange-with-Tunneling

In order to understand the mechanism of fold switching in Lymphotactin by way of computer simulations, one has to sample the free energy landscape of the protein with high accuracy. However, the accessible time scales in all-atom molecular dynamics simulations in explicit solvent are even for small proteins with less than 100 residues only of order ≈ *µs*, and therefore too small to obtain sufficient statistics. We have proposed in earlier work ^19–22^ to overcome this sampling problem in studies of conformational transitions by a variant of the Hamilton Replica Exchange method. ^23,24^ Our approach relies on two ingredients. First, a ladder of replicas is set up, where on each replica a “physical” model is coupled with a structure-based model. On one side of the ladder the structure-based model biases the physical system toward the Ltn10 state, on the other side toward Ltn40. The strength of the coupling (biasing) on each replica is controlled by a parameter λ which is maximal at the two ends, and zero for the central replica, where the physical model is therefore not biased by one of the structure-based models. Exchange moves between neighboring replicas induce a walk along the ladder by which the Lymphotactin configuration changes from one motif into the other. When these exchange moves are accepted or rejected with the criterium commonly used in Replica Exchange Sampling, the correct distribution according to the given λ-value will be sampled on each replica. Hence, on the central replica, at λ = 0, the correct and unbiased distribution of the physical model of Lymphotactin will be sampled. While formally correct, this sometimes also called Multi Scale Essential Sampling (MSES) ^25,26^ is of limited use in studies of large systems, as the acceptance probability for exchange moves becomes vanishingly small. Hence, the second ingredient of our approach is to replace the canonical acceptance criterium through a new one that allows the system to “tunnel” through the unfavorable “transition state” generated by the exchange move. This tunneling is achieved by re-scaling the velocities of atoms in the two configurations in such a way that the total energy at a given λ value is after the exchange the same as before the exchange. The two replicas evolve then by microcanonical molecular dynamics, exchanging potential and kinetic energy. After a short time (a few ps) the velocity distribution of each of the two replicas will approach the one that would be expected at the given temperature. At this point, the potential energies of the two configurations are compared with the corresponding energies before the exchange move, and either accepted or rejected. We have coined this approach as Replica-Exchange-with-Tunneling (RET), and described it and its limitations in detail in our earlier works. ^19,20^ In previous work,^20–22^ we could show that our above described approach leads indeed to an enhanced sampling of transition events and an improved statistics in the sampled energy landscape (which is generated from the unbiased central replica where λ = 0).

### 2.2 Simulation Set-up

In the present work we use our enhanced sampling method to study the conversion between Ltn10 and Ltn40 monomers. For this purpose, we simulate our system with an energy function

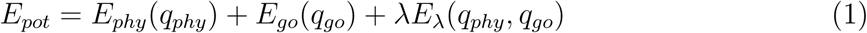

where *E*_*pot*_ is the total potential energy of the system, *E*_*phy*_(*q*_*phy*_) and *E*_*go*_(*q*_*go*_) are the potential energies of the physical model and Go-model, respectively, and *E*_λ_(*q*_*phy*_, *q*_*go*_) describes the coupling between the two models.

Interactions in the physical model are described by the CHARMM36m force field^27^ in combination with TIP3P explicit solvent,^28^ with an acetyl group cap on the N-terminus and a methylamine group cap for the C-terminus. Re-scaling the masses of the all-atom physical models by 14.49 is required to match the temperature scales of the two models as the Go-models do not include hydrogen atoms, i.e., have a smaller number of degrees of freedom. The initial configurations were randomized by high temperature molecular dynamics runs at 1500 K for 1 ns, then cooled down to the target temperature at 310 K for another 1 ns.

The two structure-based (Go-)models (one biasing toward Ltn10, the other toward Ltn40) were generated using the SMOG-Server^29^ at http://smog-server.org. While the wild-type form of Lymphotactin consists of a 93 residues, most of the C-terminus tail is disordered in both conformations, 30% for Ltn10 and 42% for Ltn40. ^10^ As the set-up of a structure-based energy function is meaningless for such unstructured regions, we have not considered this tail. Instead, we have restricted our simulations to a 75 residue fragment which describes the parts of Lymphotactin that are structured in at least one of the two folds. The length of this fragment matches the one of the NMR resolved Ltn10 form (PDB ID: 2HDM^9^), but is larger than the Ltn40 form (PDB ID: 2JP1^10^) for which only 60 residues are resolved. For the generation of the SMOG parameter the remaining 15 residues were assumed to be in a random configuration and added using the PyMOL ^30^ mutagenesis tool.

The biasing energy is defined as^25,26^

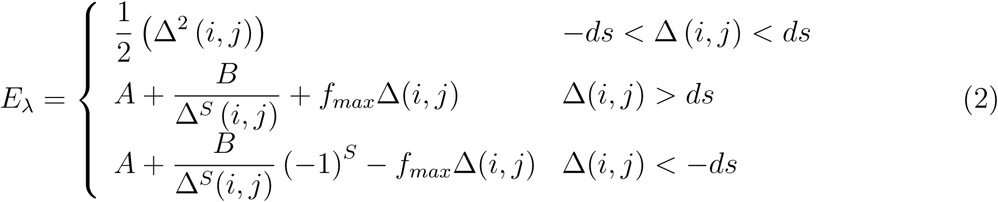

where Δ(*ij*) = *δ*_*phy*_(*ij*) − *δ*_*go*_(*ij*) is the difference of the distances between C_*α*_-atoms *i* and *j* as measured in the two models. The control parameter *f*_*max*_ sets the maximum force when Δ (*i, j*) → ∞, and *S* determines how fast this value is realized. In our case, we have *S* = 1, *f*_*max*_ = 0, and *ds* = 0.3 Å. The parameters *A* and *B* ensure continuity of *E*_λ_ and its first derivative at Δ (*i, j*) = ±*ds*, and are given by

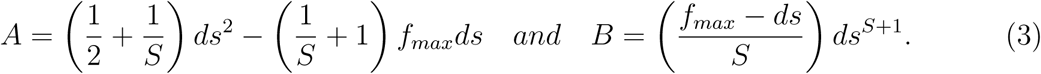

The 24 replica systems were prepared with a λ distribution of λ = 0.1, 0.09, 0.08, 0.07, 0.06, 0.05, 0.04, 0.03, 0.02, 0.01, 0.009, 0, 0, 0.009, 0.01, 0.02, 0.03, 0.04, 0.05, 0.06, 0.07, 0.08, 0.09, 0.1. Here the *E*_λ_ term biases replica 0–10 toward the Ltn10 motif, and replica 13–23 toward Ltn40. In order to simplify our programming, we use two replicas (with indices 11 and 12) to represent the case where the physical model is not biased by any structure-based biasing-term, i.e., where λ = 0.

Our simulations rely on an in-house implementation of the above described approach into the Gromacs 4.6.5 package^31^ that is available from the authors. The equations of motion are integrated with the Velocity Verlet algorithm, ^32^ with hydrogen bonds constrained by the LINCS algorithm,^33^ using a time step of 2 fs. The van der Waals and electrostatic cutoffs are set to 1.2 nm. Note that instead of simulating each of the replicas at a constant temperature, the temperature of the replicas is changed in steps of 0.01 K between 310 and 310.23 K due to the way our code has been implemented into Gromacs, where the temperature of the replicas is maintained by using the v-rescale thermostat.^34^ Choosing the length of the microcanonical segment in the RET move as 1 ps, a 100 ns trajectory was generated. Thermodynamic quantities were calculated solely from replica with λ = 0, i.e., such without bias from the two structure-based models.

### 2.3 Observables

The free energy as a function of order parameter, λ, is defined as

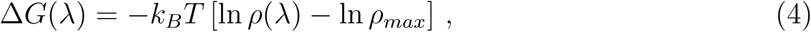

where *k*_*B*_ and *T* denote the Boltzmann’s constant and temperature, respectively. *ρ* is an estimate of the probability density function calculated from a histogram of the data, while *ρ*_*max*_ is the maximum of the density. The second term ensures that Δ*G* = 0 for the lowest free energy minimum. Free energy values reported in this work are calculated at a temperature of 310 K.

The configurations as obtained from the equilibrated trajectories corresponding to unbiased trajectories (λ = 0) are characterized based on secondary structure pattern calculated using STRIDE algorithm^35^ as implemented in VMD software^36^ and latter discussed characteristic backbone hydrogen bonding pattern specific to each of Ltn native forms (see Table 1). Ltn has four characteristic *β*-strands, *β*_0_ (residue 10–15), *β*_1_ (residue 25–31), *β*_2_ (residue 34–41), *β*_3_ (residue 44–51) and a C-terminal helix, H (residue 54–66). Based on the secondary structural pattern of these five regions and backbone hydrogen bonding pattern in *β*_1_ −*β*_2_ −*β*_3_ region, we define three variables (B0, H, and B123) using following logical expression, which eventually helps us characterize the configurations and distinguish the pattern among them.

**Table 1:** Characteristic Backbone Hydrogen Bonding Pattern among Different *β*-Strands for Ltn10 and Ltn40 Lymphotactin Native Forms as Obtained from Our Study and Reported in the NMR Structure.^10^

| $\beta_1 \leftrightarrow \beta_2$ | | $\beta_2 \leftrightarrow \beta_3$ | |
| --- | --- | --- | --- |
| Ltn10 | Ltn40 | Ltn10 | Ltn40 |
| T26 ↔ I40 | K25 ↔ I40 | – | R35 ↔ D50 |
| T28 ↔ I38 | Y27 ↔ I38 | V37 ↔ A49 | V37 ↔ C48 |
| T30 ↔ A36 | I29 ↔ A36 | F39 ↔ V47 | F39 ↔ K46 |
| – | E31 ↔ L34 | T41 ↔ L45 | T41 ↔ G44 |

- **B**0: If there are at least two residues in *β*-strand in *β*_0_ region, then the value of B0 is 1 i.e., *β*_0_ exists; otherwise it is zero.
- **H**: If there are at least three residues in helix in C-terminal helix region, then the value of H is 1 i.e., C-terminal helix exists; otherwise it is zero.
- **B**123: If there is no characteristic hydrogen bonds (Ltn10-like or Ltn40-like) in *β*_1_ − *β*_2_ −*β*_3_ region and there is less than two residues in *β*-strand in *β*_1_ −*β*_2_ or *β*_2_ −*β*_3_ regions, B123 will be assigned to have a value of zero. If this condition is not satisfied, B123 will have hydrogen bonding pattern as that of Ltn10-like or Ltn40-like or Mixed (i.e., substantial existence of both type of hydrogen bonds) depending on the conditions: if the difference in number of characteristic Ltn10-like and Ltn40-like hydrogen bonds (see Table 1) is greater than or equal to two, B123 is Ltn10-like. Similarly, if the difference in number of characteristic Ltn40-like and Ltn10-like hydrogen bonds is greater than or equal to two, B123 is Ltn40-like. All other remaining configurations will be considered to have mixed hydrogen bonding pattern.

One quantity by which we have measured similarity of a given configuration to one of the two native Lymphotactin folds is the fraction of specific native contacts *Q*_*spec*,1_(*X*), defined as

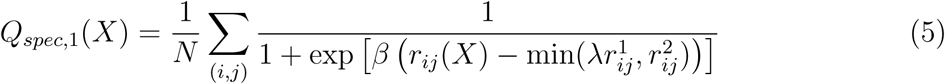

Here, 1 denotes one native fold of Lymphotactin (Ltn10 or Ltn40), while 2 represents the alternative one. Only contacts specific to either of the native folds are considered, i.e., contacts that are found in both native folds are excluded. Here, we define a contact in the native structure by the requirement that two backbone atoms on distinct residues are within 4.5 Å. Thus, *N* is the number of such contact pairs (specific to the one form of Lymphotactin native structure) of (*i, j*) backbone atoms *i* and *j* belonging to residues *θ*_*i*_ and *θ*_*j*_. To avoid the contacts forming by adjacent residues, we have considered only the residues where | *θ*_*i*_ − *θ*_*j*_ |> 3. *r*_*ij*_(*X*) is the distance between the atoms *i* and *j* in conformation *X*, while 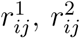 are that distance in the native fold 1 and 2, respectively. *β* is used to smooth the distribution of the values and considered to be 5 Å^−1^. The fluctuation of the contact formation is controlled by the minimum value of λ times 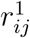 and 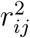, where λ is taken to be 1.8. By introducing such a minimum value between the above mentioned two quantities, we set a range of fluctuation for each of the specific native contacts considered so that one could differentiate between the set of specific native contacts with respect to the two folds, and thus measure the similarity of a given configuration with respect to only one of the two native folds.

## 3 RESULTS AND DISCUSSION

Previous investigations into the conversion mechanism between the two Lymphotactin motifs were hampered by the computational difficulties of sampling the protein’s free energy landscape with sufficient statistics. In regular molecular dynamics simulations, the protein will spend most of the time in one of the basins of attraction - either exploring configurations similar to Ltn10, or, in the other case, Ltn40-like configurations - with transitions between the two basins being rare events. We argue that these computational difficulties can be overcome with our variant of replica exchange sampling which was designed specifically for investigation of switching between two well-defined states. In order to demonstrate that our approach allows indeed a more accurate sampling of the free energy landscape by raising the rate of transitions between the two main basins, we show in Figure 2(a) the walk of a typical realization of our system along the ladder of replicas. At start time (*t* = 0) the physical system sits on a replica where it is biased with λ = 0.009 toward the Ltn40 form. During the 100 ns of simulation this realization of Lymphotactin walks numerous times between the two end-points of the ladder. On the one end, the physical system will be biased maximally with coupling parameter λ_*max*_ = 0.1 toward the Ltn40 structure, and on the other end with a maximal λ_*max*_ = 0.1 toward Ltn10. The average exchange rate between neighboring replicas is ∼ 47%. In Figure 2(a) we show that this walk through λ-space induces indeed interconversion between the two forms. For this purpose, we characterize the state of a given configuration by the C_*α*_ distances between two specific residue pairs. In the Ltn10 structure the two residues T15 and A49 form a hydrophobic contact, while they are separated by a large distance in Ltn40. The opposite relation is found for the pair L14 and L45, which form hydrophobic contacts in Ltn40 but are far away from each other in the Ltn10 structure. Both distances are shown in Figure 2(b) and evolve in anti-correlated fashion over the 100 ns of simulation. If the system is on one side of the ladder and biased toward Ltn10, the distance between T15 and A49 (shown in red) will have small values and the distance between L14 and L45 (drawn in black) have large values, while the opposite is true once the system is on replica where the bias from the structure-based model is toward the Ltn40 structure.

**Figure 2:**
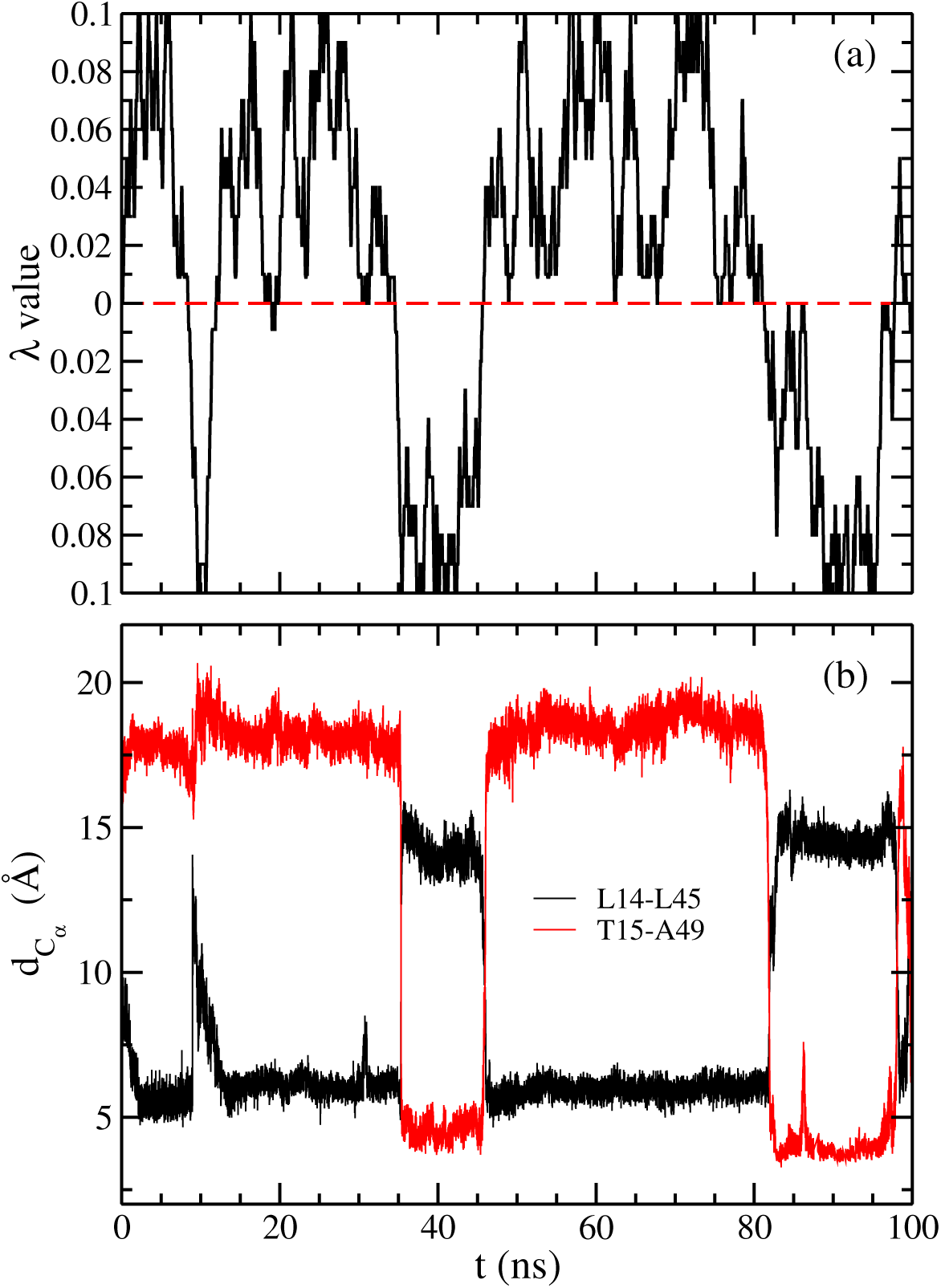
(a) A typical example of replica walking through λ space starting from a replica, where the physical model is initially biased toward Ltn40 with λ = 0.009. While the system walks between replicas with bias toward Ltn10 and such with bias toward Ltn40, its configuration changes accordingly. This can be seen in (b) where we show the corresponding time evolution of C_*α*_ distances 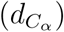 between L14 and L45 (black) and that between T15 and A49 (red). The first distance is a measure for the similarity with Ltn40, and the second distance one for the similarity with Ltn10.

A measure for the efficiency of our method, and a lower limit on the number of independent configurations sampled at the λ = 0 replicas, is the number of walks along the whole ladder, from the replica with maximal bias toward Ltn10 to the one with maximal bias toward Ltn40, and back. Such a walk along the whole ladder of replicas is called by us a tunneling event, and inversely related to the average time needed to cross the ladder (termed by us the tunneling time). The higher the number of tunneling events, and the shorter the tunneling time, the more efficient will be our approach. However, calculation of the number of tunneling events or the tunneling time gives only meaningful results after the system has approached equilibrium. This convergence of the simulation is checked by calculating the free energy as a function of the two distances introduced above, and comparing it for different time intervals. Resemblance of the data for different intervals in Figure 3 suggests that the simulation has converged after 20 ns, and therefore, we use the last 80 ns of the simulated trajectories for our analysis. In this time span, we find a total of 31 tunneling events with an average tunneling time of ∼ 33 ns. We have demonstrated in earlier work^19–22^ for smaller systems that the number of tunneling events is in our approach much higher than the ones found in regular Hamilton replica exchange simulations with comparable number of replicas and λ distribution. In the latter case, often not a single tunneling event could be detected. As Lymphotactin is larger than the previously studied systems, and the sampling difficulties increase exponentially with system size, we expect that for Lymphotactin the improvement over regular Hamilton replica exchange is even higher than seen in our previous work.

**Figure 3:**
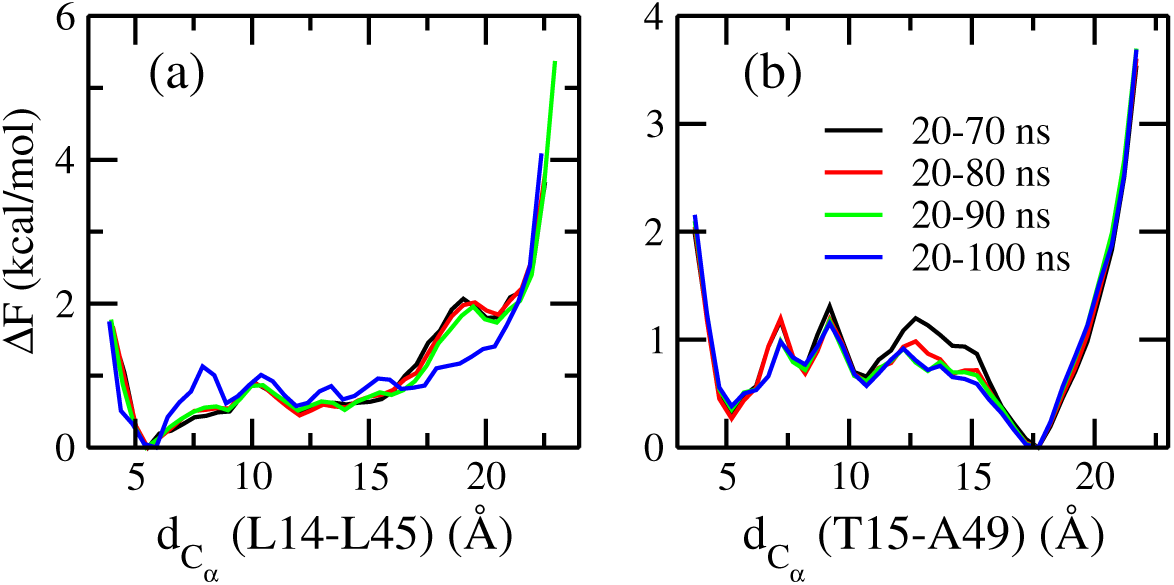
Free energy (Δ*F*) as a function of the two C_*α*_ distances 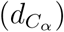 between (a) L14 and L45 (a measure for the similarity to Ltn40) and (b) that between T15 and A49 (a measure for the similarity to Ltn10), as obtained from the unbiased replica where λ = 0. Shown are values as measured for different segments of the simulation.

The increased efficiency of our approach, leading to 31 tunneling events, gives us confidence in the free energy landscape found at λ = 0, i.e., at a replica where the “physical” model of our system is not biased toward either Ltn10 or Ltn40. In order to measure the frequency of these two motifs for the unbiased replica, we define a configuration as Ltn10-like if the C_*α*_ distances between T15 and A49, 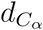 (T15-A49), is less than 8 Å and and that between L14 and L45, 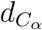 (L14-L45), greater than 12 Å. This definition was derived from visual inspection of the landscape as obtained for the replica with maximal bias toward Ltn10. Guided in a similar way by a visual inspection of the landscape as obtained for the replica with maximal bias toward Ltn40, we define a configuration as Ltn40-like if 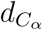 (L14-L45) is less than 8 Å and 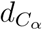 (T15-A49) greater than 12 Å. Using the above definitions we find that on the unbiased replica about 17% of configurations are Ltn10-like, while 34% are Ltn40-like. This suggests that Ltn40 is the most stable form and about 50% of the configurations sampled in our simulation do not represent either fold. Visual inspection and secondary structure analysis indicate that most of those configurations are intermediates on the pathway between the two opposite folds. The slightly higher population of Ltn40 over Ltn10 is in accord with the experimental study, ^15^ where under physiological conditions 46% of the configurations are Ltn10-like, and 54% Ltn40-like. On the other hand, configurations that do not belong to either of the two motifs are not observed with the large frequency seen in our simulations. This difference can be explained by the low temporal resolution of the experiments which makes it difficult to characterize short-lived intermediates, i.e., the experimentally reported frequencies for Ltn10 and Ltn40-like configurations are relative frequencies resulting from the well-resolved signals corresponding to 12 residues collected from a two-dimensional ^1^5N-^1^H HSQC spectrum.

In order to explore the interconversion pathway, we show in Figure 4(a) the free energy landscape projected on the two characteristic distances introduced earlier: 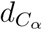 (L14-L45) (measuring similarity with Ltn40) and 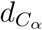 (T15-A49) (quantifying similarity to Ltn10). For calculating the landscape, we use only data sampled at the λ = 0 replicas, where the physical model is not biased by any Go-model term. The so-drawn landscape is characterized by two prominent basins corresponding to either Ltn10-like or Ltn40-like configurations. Conversion events can be described by pathways connecting the two basins in the landscape. However, not all possible pathways are equally likely. Take as an example the pathway represented by a black line in Figure 4(a). This path is obtained by projecting onto the landscape the configurations sampled along the walk in λ space during a certain tunneling event. This pathway describes a transition between the two Lymphotactin folds that requires crossing an energy barrier much higher than that reported in the experiments, indicating that this is not true pathway. On the other hand, we can obtain a thermodynamically reasonable pathway by using the MEPSA software^37^ to construct the minimum energy pathway, which we have drawn as a white line in the landscape in Figure 4(a). Unlike the tunneling pathway, this pathway does not go through regions of the landscape characterized by unfolded configurations, but instead proceeds through a series of basins, with an effective energy barrier similar to that reported in experiments. This indicates that the interconversion process does not proceed by unfolding of the Ltn10 or Ltn40 structure but rather involves a series of intermediates or transition states

**Figure 4:**
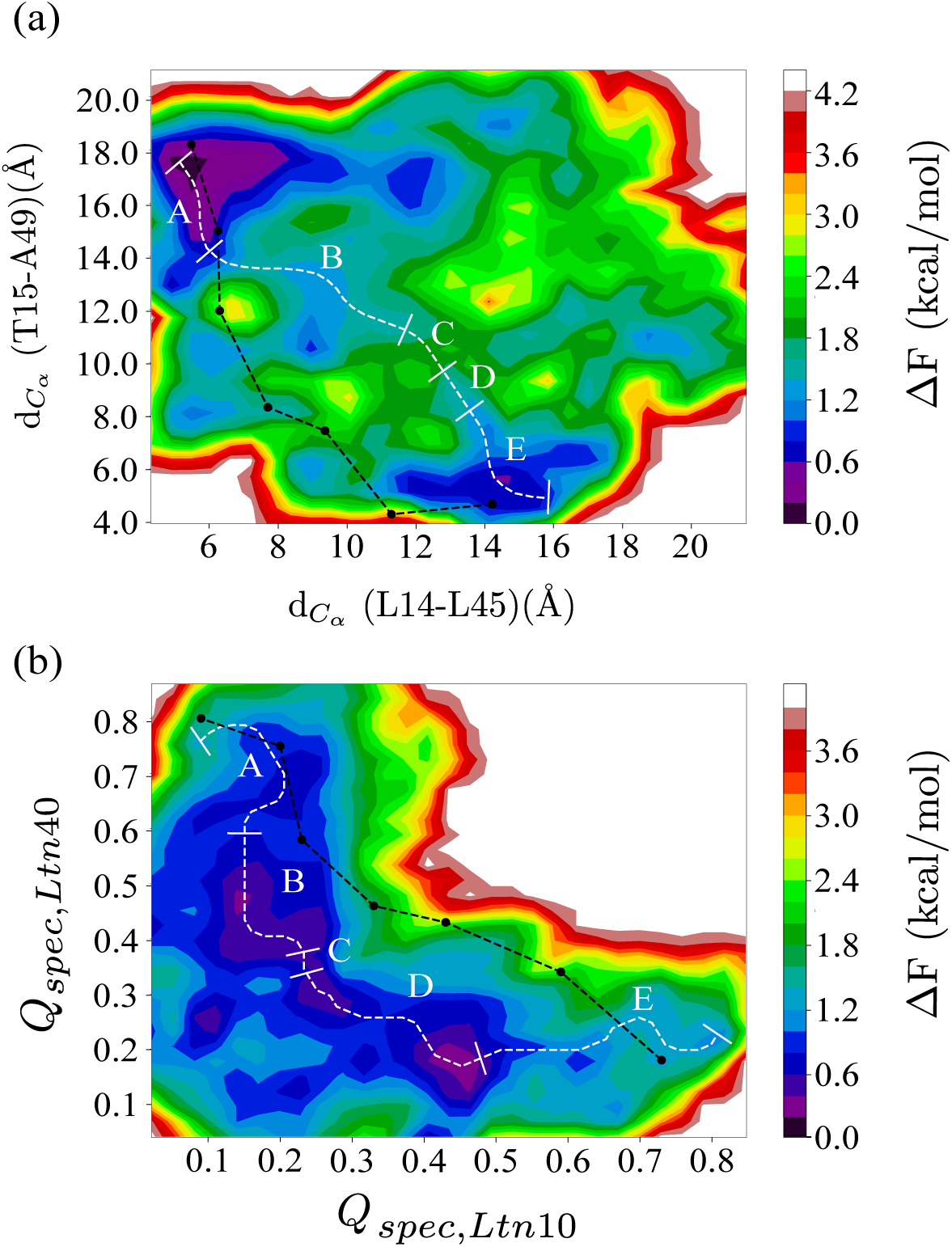
Free energy landscape as obtained from our RET simulation at replicas where the physical models are not biased by any Go-term. The landscape is in (a) projected on C_*α*_ distances between L14 and L45 ((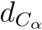 (L14-L45)), a measure for similarity to the Ltn40 structure) and that between T15 and A49 ((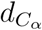 (T15-A49), a measure for the similarity to Ltn10). The black line shows a typical pathway as obtained during a tunneling event, i.e, from a walk in λ-space between the structure strongly biasing toward Ltn10 and that toward Ltn40. On the other side, the minimum energy pathway as obtained from MEPSA software^37^ is drawn in white. The labels A-E mark five distinct regions that can be identified along this pathway (discussed in the text). For comparison, we show in (b) the same landscape, but projected on the fraction of specific native contacts (defined in Eqn. 5) with respect to Ltn10 (*Q*_*spec,Ltn*10_) and Ltn40 (*Q*_*spec,Ltn*40_) native structure. Both the minimum energy and the tunneling pathways are shown again, using the same color coding as in (a).

By construction the minimum energy pathway does not connect specific configurations but bins in the landscape each containing a certain number of configurations sampled throughout the simulations. These configurations can be characterized according to presence (or lack) of a C-terminal helix (found in the Ltn10 structure), the N-germinal *β*_0_ strand (found in the Ltn40-structure), and the hydrogen bonding in the *β*_1_ to *β*_3_ region, which offers another way of distinguishing between the Ltn10 and Ltn40-like configurations. The procedure by which we attribute these three traits to a configuration is described in the method section. The frequency with which the three traits are observed allows one to identify five distinct regions along the pathway that correlate with the basins and barriers of the landscape. These segments are labeled as A to E in Figure 4(a). A similar division is not possible for the pathway, derived from the tunneling event

The difference between the two possible pathways does not depend on the specific coordinates on which the landscape is projected. This can be seen from Figure 4(b) where we project the free energy landscape on the fraction of specific native contacts with respect to each of the two folds, and overlay again both paths on the landscape. The pathway derived from the tunneling event is again not consistent with the landscape. Hence, the tunneling events in our RET approach cannot be used to derive a conversion mechanism as they rely on an artificial dynamics, designed to increase sampling efficiency. On the other hand, while the configurations in the minimum energy pathway calculated for the new landscape will differ from the one calculated for the other landscape, we can again identify the same five regions. We remark that the radius of gyration (a measure for the compactness of configurations) of the central part, made up of *β*_1_ − *β*_2_ − *β*_3_ in both Ltn10 and Ltn40, differs little along the pathway, which implies that the Lymphotactin configurations do not unfold and refold while assuming this pathway during the interconversion process.

The qualitative agreement between the minimum energy pathways found for the two landscapes suggests that these pathways describe indeed the conversion process. Hence, in order to determine the mechanism and to establish the separating free energy barrier, we have analyzed in more detail the pathway shown in Figure 4(a). When starting from the Ltn40 basin of region A, the *β*_0_ strand gets dissolved when reaching basin B which has about 1.5 kcal/mol higher free energy than the Ltn40 fold of basin A. Upon crossing this barrier, the Lymphotactin configuration evolves further through a number of intermediates with little difference in free energy, forming the flat-floored valley of region C. Using visual inspection and the STRIDE algorithm^35^ as implemented in VMD software^36^ for secondary structure analysis, we observe a re-arrangement of backbone hydrogen bonds, until, when entering region D the hydrogen bond pattern of the remaining three *β*-strands (*β*_1_, *β*_2_, and *β*_3_) becomes similar to that of the Ltn10 fold. The Ltn10 basin (region E) has again about 1.5 kcal/mol lower free energy value than the transition state D and is reached when the helix slowly begins to form at the C-terminus (within the residues 54 to 66). A schematic diagram explaining the mechanism is shown in Figure 5.

**Figure 5:**
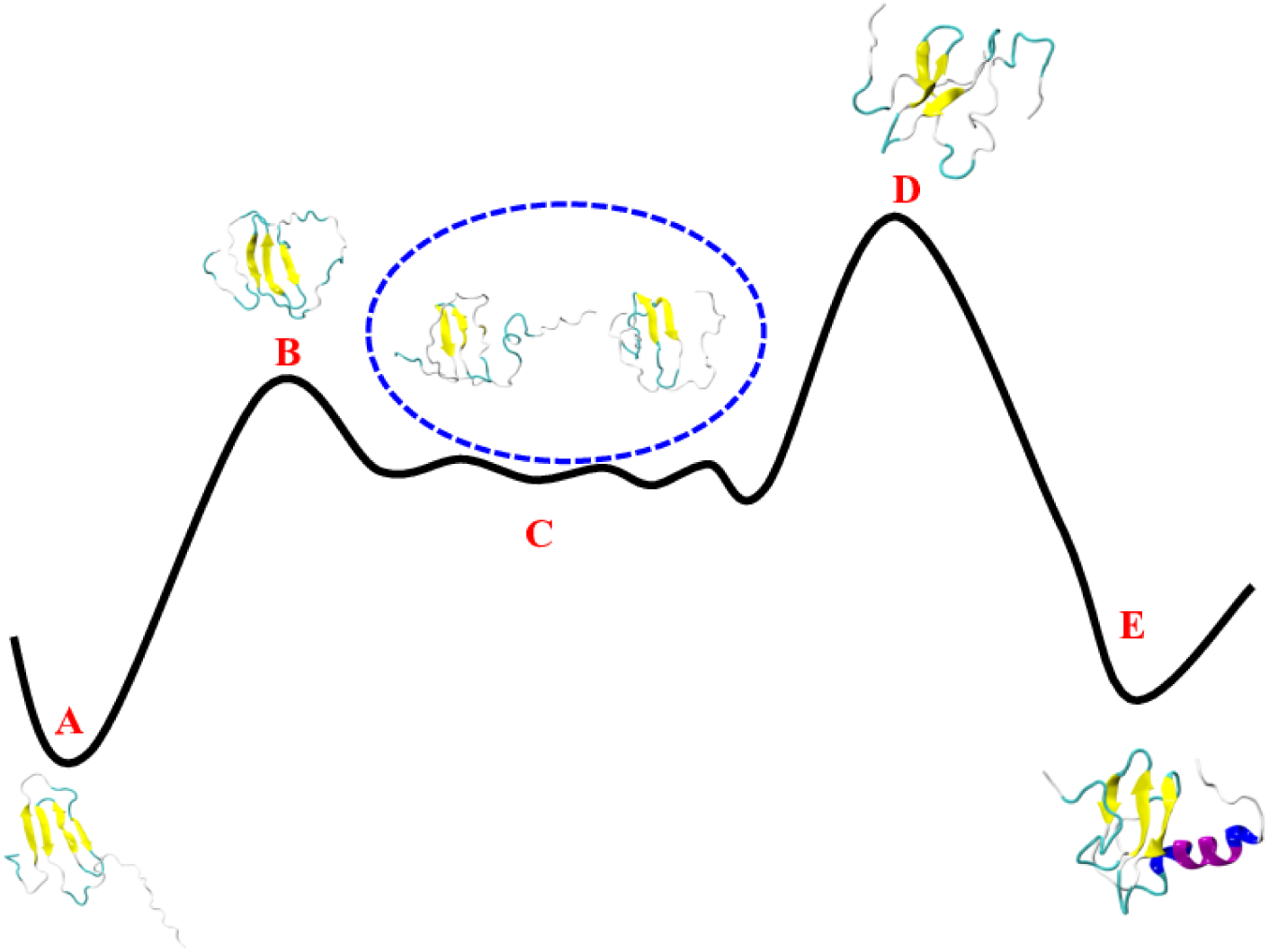
A schematic diagram of the interconversion mechanism between Ltn40 and Ltn10 with representative structures.

From the above discussed conversion process, we have estimated a total energy barrier of ∼ 2 kcal/mol for Ltn40→Ltn10 conversion, while slightly a lower value of ∼ 1.5 kcal/mol is the barrier for the reverse process. While the higher energy barrier for the Ltn40→Ltn10 conversion, implying a slower conversion rate than the reverse one, is in agreement with the experimentally reported results, ^11,17,38^ the energies are higher than the ones measured in the experiments: 0.5 kcal/mol and 0.2 kcal/mol, respectively. This divergence may reflect again the difference between our computational setting (considering a transition between monomer structures) and the experimental setting that studies the transition from Ltn10 monomers to Ltn40 dimers. As the dimerization is the fast process, the Le Chatelier’s principle will predict a shift of the equilibrium toward the Ltn40 motif, effectively lowering the barrier.

Unlike an earlier proposal, ^17^ our conversion mechanism does not require unfolding of the Ltn monomers. Instead it relies on a sequence of local changes with the main assumption that the intermediates keep part of the local ordering. Especially, we assume that the central three-stranded *β*-sheet (*β*_1_, *β*_2_, and *β*_3_) is preserved. This sheet is found in both motifs, but with a shift in the backbone hydrogen pattern by one residue,^10^ see Table 1. Note that residues on the central *β*_2_ strand do not get shifted. Instead, they change their hydrogen bond forming partners on the adjacent strands (*β*_1_ or *β*_3_). Our proposed conversion mechanism therefore has to explain how this shift between these residues can proceed without the need of a complete unfolding. A more thorough analysis of the pathway shows that the unfolding of the *β*-sheet and the resulting high barriers are avoided by some residues forming bifurcated/bridged hydrogen bonds with two consecutive residues. For example, while in the Ltn40 configurations of basin A, residue F39 forms a backbone hydrogen bond with residue K46, and in the Ltn10 configurations of basin E backbone hydrogen bonds with residue V47. However, we observe that F39 (located on strand *β*_2_) can also form simultaneously hydrogen bonds with residues K46 and V47 on strand *β*_3_. In this case, the amide nitrogen of F39 only participates in forming hydrogen bonds and thus acts as a bifurcated donor, or both carbonyl oxygen and amide nitrogen of F39 participate in forming those hydrogen bonds, helping in forming bridged hydrogen bonds. Similarly, we find that in a similar manner residue T41 can also form hydrogen bonds with both residues G44 and L45. All six residues on the central *β*_2_ strand (see Table 1) can form such bifurcated hydrogen bonds that bridge between Ltn10-like and Ltn40-like hydrogen bonding, but for most residues these bifurcated hydrogen bonds appear with a relative frequency of less than 5%, that is, less than 5% of all hydrogen bonds connecting a residue on strand *β*_2_ with a partner residue on either *β*_1_ or *β*_3_ are bifurcated hydrogen bonds.

The exceptions are the already mentioned bifurcated hydrogen bonds F39-K46/V47, which is observed with a relative frequency of about 24%, and T41-G44/L45, which appears with a relative frequency of about 11%. We remark that we find similar numbers also for the minimum free energy path of the free energy landscape in Figure 4(b). Both bifurcated hydrogen bonds connect residues located at the start of the sheet formed by the *β*_2_ and *β*_3_ strand. In the Ltn10 motif, residue F39 forms a hydrogen bond with V47, and T41 one with L45. Transitioning to the Ltn40 hydrogen bonding of F39 with K46 and T41 with G44 will be eased by transient formation of the bifurcated hydrogen bonds at this location as it avoids the energetic costs of dissolving and reforming hydrogen bonds. Correspondingly, the F39-K46/V47 is observed with highest relative frequency in the transition regions B (about 37%) and D (about 64%), and with about 16% relative frequency in the intermediate region C. The corresponding frequencies are lower for the T41-G44/L45 bifurcated hydrogen bond, which appears in 8% (11%) in the transition region B (D), and with a relative frequency of 14% in the intermediate region C.

We conjecture that formation of the bifurcated/bridged hydrogen bonds F39-K46/V47 and T41-G44/L45 at the turn region between the *β*_2_ and *β*_3_ strands is crucial for enabling the fold switch as it disturbs the geometry of the sheet and initiate a wave of successive rearrangement hydrogen bonds in the three-stranded *β*-sheet which avoids large energy barrier that would otherwise arise from breaking and forming hydrogen bonds. Our conjecture could be tested in principle by mutation experiments where the mutated side chain would form contacts that lead to repositioning of the backbone atoms that would restrict formation of such bifurcated hydrogen bonds. Another possibility would be the use of deuterium which would also alter the relative frequency of bifurcated hydrogen bonds.

Interestingly, both bifurcated hydrogen bond pairs are also seen with substantial frequency in the region E (dominated by Ltn10-like configurations), where the F39-K46/V47 bond is observed in about 49% of the configurations, and the T41-G44/L45 one in about 16% of configurations. Hence, in Ltn10-like configurations the hydrogen bond pairs F39-V47 and T41-L45 are easily replaced by the corresponding bifurcated hydrogen bonds. This is likely because the extension from the hydrogen bond F39-V47 to the bifurcated hydrogen bond F39-K46/V47 involves rotation of the side chains of the three residues that reduces the hydrophobic solvent accessible surface area (SASA) by about 26 Å^2^, and increases the exposure of the charged K46 by about 7 Å^2^. Both are energetically favorable changes. The effects are less pronounced for the bifurcated hydrogen bond T41-G44/L45 where there is only a small reduction in hydrophobic SASA ≈ 7 Å^2^. On the other hand, in region A (where Ltn40-like configurations are found) only the T41-G44/L45 bifurcated hydrogen bond is observed with substantial, but much smaller, relative frequency (about 9%). Here, formation of a bifurcated hydrogen bond T41-G44/L45 would not change the solvent accessible surface area of the three involved residues, while the bifurcated hydrogen bond F39-K46/V47 would lead to a reduction of about 45 Å^2^ of solvent exposure for the charged K46 that could not be compensated by the favorable loss of hydrophobic SASA of about 18 Å^2^. However, we remark that unlike to the Ltn10 configuration, where the *β*_3_ strand is stabilized by four contacts with the N-terminal helix, no such *β*_3_-stabilizing contacts with the *β*_0_ strand exist in the Ltn40 configuration. Hence, the barrier for unraveling the *β*_2_ − *β*_3_ hydrogen bonding is likely lower in the Ltn40 configurations than in the Ltn10 configurations where instead formation of the two bifurcated hydrogen bonds circumvents the otherwise higher barrier.

Differences in the number of stabilizing side chain contacts are also the reason why we observe bifurcated hydrogen bonds with appreciable frequency only between *β*_2_ and *β*_3_, but not between *β*_2_ and *β*_1_. In the Ltn10 configuration, four side chain contacts (A49-W55, A49-V56, D50-V56, P51-V56) with the helix stabilize the *β*_3_ strand, but ten such side chain contacts (L24-W55, L24-C59, L24-M63, L24-K66, T26-M63, Y27-V60, Y27-R61, Y27-M63, Y27-D64, I29-V60) that stabilize the *β*_1_ strand. The difference in number of stabilizing side chain contacts is even larger for the Ltn40 configuration where there are no side chain contacts with the *β*_0_ strand that could stabilize the *β*_3_ strand, but for such contacts that stabilize the *β*_1_ strand. As formation of bifurcated hydrogen bonds requires repositioning of backbone atoms that in turn depends on suitable rotation of side chain atoms, such bifurcated hydrogen bonds are less likely between *β*_1_ and *β*_2_ than between *β*_2_ and the more flexible *β*_3_.

## 4 CONCLUSIONS

Using a variant of Replica-Exchange-with-Tunneling (RET), we have probed the interconversion of a metamorphic protein that switches biological function by altering its three dimensional structure between two well-defined native forms, Ltn10 and Ltn40. While Ltn10 is monomer with three-stranded *β*-sheet ending with a C-terminal helix, Ltn40 exists as a dimer with all *β*-sheet arrangements. In order to ease the numerical difficulties, we have only considered the conversion for a monomer. This is justified as conversion and dimerization are separate events, with the conversion the time-limiting process. ^17^ Our investigation relies on the use of RET, an enhanced sampling method developed in our group, that has enabled us to sample the free energy landscape of the protein with high precision. We find relative population frequencies that are consistent with experimental measurements, but our simulations predict a larger population of intermediate configurations than reported in the experiments. We reason that our method allows us to identify intermediates that due to their short-life time are difficult to observe in experiments. Analyzing the free energy landscape allows us to identify an conversion mechanism that relies on passage through a number of distinct structural intermediates, and involves breaking and reformation of the *β*_0_-strand and the C-terminal helix and a re-arrangement of hydrogen bonds in the central three-stranded *β*-sheet made from *β*_1_, *β*_2_ and *β*_3_. The associated high costs of breaking and forming hydrogen bonds are avoided by formation of bifurcated hydrogen bonds that naturally bridge between the characteristic hydrogen bond pattern in the three *β*-sheets common in both motifs. These pattern differs in both forms by being shifted by one residue. We surmise that formation of these bifurcated hydrogen bonds facilitates the switch between these two patterns, guiding in this way the conversion between the two motifs.

## Acknowledgement

The simulations in this work were done using the SCHOONER cluster of the University of Oklahoma and XSEDE resources allocated under grant MCB160005 (National Science Foundation). We acknowledge financial support from the National Institutes of Health under grant GM120634.

